# Effects of Alternating Polarity Frequency on the Assembly of Electro-Methanogenic Microbial Communities

**DOI:** 10.64898/2026.10.16.759672

**Authors:** Yu Liu, Shiyun Yao, Weibo Xia, Bingxin Li, Xinghan Zhao, Qiang He, Heyang Yuan

## Abstract

Electro-methanogenesis can enhance methane production and has attracted extensive interest, but the current strategies for building electro-methanogenic microbial communities suffer from several limitations. Alternating polarity has been shown as a new strategy to build electro-methanogenic communities, but the impact of alternation frequency on community assembly remains to be understood. In this study, the impact was evaluated using bioelectrochemical systems amended with granular activated carbon, and the working-electrode potential was alternated between +0.4 and -0.4 V vs. SHE at six frequencies ranging from 0.5 to 12 days. Among the tested conditions, 4-d alternation led to the best overall performance, producing approximately 250 L/L/kg cumulative biogas, 96% methane content, and a 45% methane yield, while preserving a stable and reproducible community structure. The community contained two *Methanobacterium* populations, two putatively electroactive *Geobacter* populations, and one putatively electroactive *Pseudomonas* population, together with diverse fermentative and syntrophic bacteria. Transcript-based analyses revealed clear frequency- and potential-dependent shifts in the activity of these functional groups. Redundancy analysis associated the *Methanobacterium formicicum*-related population with cathodic current and methane production, whereas the *Geobacter sulfurreducens*-related population was linked to anodic current and methane production. Bayesian and local-similarity networks further connected methanogen dynamics with specific electroactive, fermentative, and syntrophic partners. Collectively, these results establish alternating polarity frequency as a critical parameter for building robust communities capable of high-rate, high-quality methane production.

## 1. Introduction

Methanogenesis is a ubiquitous microbial process found in natural and engineered systems.^1^ In engineered systems such as anaerobic digesters, methanogenesis is coupled with upstream processes including hydrolysis, acidogenesis, and acetogenesis to convert organic wastes into methane-rich biogas, thereby providing a sustainable solution to simultaneously treat waste streams and produce energy.^2–4^ Despite the great promise of methanogenesis in practical applications, numerous studies have highlighted its critical limitations. In particular, classical methanogenesis via the hydrogenotrophic, acetolactic and methylotrophic pathways is a slow process limited by the activity of upstream microbial processes.^5, 6^ There is continuous interest in developing new methods to overcome the low kinetics of classical methanogenesis.^7, 8^

Electro-methanogenesis as a new methanogenic pathway has been reported to produce methane-rich biogas at high rates. In this process, electrotrophic methanogens accept extracellular electrons from electrodes and/or electroactive partners such as *Geobacter* to reduce carbon dioxide (CO_2_) to methane.^9^ Electro-methanogenesis was first demonstrated using electrodes as the electron donor,^10^ and later studies reported high efficiencies of electron capture (>80%) and CO_2_-to-CH_4_ conversion (98%), as well as a high methane production rate of 0.055 mol/day/mg-biomass with the cathode potential more negative than -0.6 V vs. standard hydrogen electrode (SHE).^11, 12^ In addition, electro-methanogenesis can also happen via direct interspecies electron transfer (DIET), as shown by early studies in which *Geobacter metallireducens* directly transferred electrons to *Methanosarcina barkeri*.^13^ Mediated by conductive materials such as granular activated carbon (GAC), anthracite, and biochar, DIET between electrotrophic methanogens and electroactive partners has been reported to achieve 86% electron recovery from ethanol to methane and increase the specific methane yield by 43%.^14–16^ As the ability of both electrode- and DIET-driven electro-methanogenesis to enhance biogas production and methane content has been consistent confirmed,^17–20^ there is a continuous interest to develop strategies to build electro-methanogenic microbial communities.

Three strategies have been shown to promote electro-methanogenesis: cathodic enrichment, microbial electrolysis, and conductive material amendment. A cathode can facilitate the enrichment of electrotrophic methanogens by acting as a stable electron source. However, cathodic enrichment-assembled communities are susceptible to operational perturbations, such as temporary cathode disconnection, in which case electrotrophic methanogens lack alternative electron sources and can no longer perform electro-methanogenesis.^21, 22^ Microbial electrolysis suffer from the same issue, as the electrotrophic methanogens enriched on the cathode are physically separated from the electroactive bacteria enriched on the anode and cannot use them as the alternative electron source.^23–25^ Moreover, the cathodic potential during microbial electrolysis is not precisely controlled and often more negative than -0.414 V vs. SHE, leading to abiotic hydrogen evolution that favors hydrogenotrophic methanogenesis.^26, 27^ Compared to those strategies, conductive material amendment can facilitate DIET but do not impose a strong selective pressure for enriching electro-methanogenic populations.^28, 29^ For instance, the electroactive partner, *Geobacter*, can readily divert electrons to other electroactive bacteria or involve in syntrophic growth with hydrogenotrophic methanogens.^30^ Collectively, the three existing strategies each suffers from critical issues,^31^ and new strategies are needed to enrich both electrotrophic methanogens and their electroactive partners dedicated to electro-methanogenesis.^32^

Alternating polarity has emerged as a promising strategy for assembling electro-methanogenic communities.^33, 34^ During alternating polarity, the electrode potential is periodically switched between anodic and cathodic states to impose selective pressures. This allows the electrode to alternately function as an electron donor for electrotrophic methanogens and an electron acceptor for electroactive bacteria, resulting in simultaneous enrichment of both populations.^35–37^ In the presence of a cathode, electroactive bacteria can supply additional electrons to enhance electro-methanogenesis, whereas in the absence of a cathode, they function as alternative electron donors that help sustain electro-methanogenic activity.^38, 39^ The feasibility of alternating polarity to build electro-methanogenic communities was recently confirmed.^33^ In two-chamber bioelectrochemical systems supplemented with GAC, the electrode potential was periodically alternated between +0.8 V and -0.4 V vs. SHE, with an three-day open circuit cycle between each alternation to avoid electrical shock. Under alternating polarity, cumulative biogas production increased to 125 L/L/kg-GAC after four enrichments and outperformed other non-alternating conditions. Community analysis revealed enrichment and upregulated activity of putatively electrotrophic *Methanobacterium* and electroactive *Geobacter* under cathodic and anodic potentials, respectively.^33^ Network analysis further inferred the potential syntrophic association between these populations.^33^

Despite its promise in assembling electro-methanogenic communities, the operating conditions of alternating polarity, particularly the alternation frequency, remain to be optimized. The alternation frequency can play a critical role in shaping the structure of electro-methanogenic communities, as the enrichment of electrotrophic methanogens and their electroactive partner is largely determined by the duration of the cathodic and anodic potential.^40^ Limited evidence has shown the adverse effect of overly frequent shift between cathodic and anodic potential (e.g., 10 min or 2 h) on microbial metabolic processes and growth.^41^ Such rapid perturbations can destabilize biofilm structure and impair the establishment of efficient electron transfer pathways, preventing both methanogens and electroactive bacteria from gaining dominance under the changing redox condition.^42^ Conversely, when the alternation frequency is too low, the applied cathodic or anodic potentials may be unable to supply or withdraw sufficient electrons, potentially leading to unbalanced enrichment of either electrotrophic methanogens or electroactive bacteria.^33^ It is reasonable to hypothesize that the alternation frequency should be approximately the doubling time of the populations to be enriched (e.g. 0.1-12.4 days for methanogens) so that they can complete one full growth cycle under a specific redox condition.^43–45^

The goal of this study was to systematically evaluate the effect of the frequency of alternating polarity on the assembly of electro-methanogenic communities. The specific objectives include: 1) identifying the optimal alternation frequency for rapid biogas production and high methane content, 2) characterizing the electro-methanogenic communities assembled under different alternation frequencies, and 3) understanding the activity and interactions of electrotrophic methanogens and their electroactive partners under different alternation frequencies. To this end, two-chamber bioelectrochemical systems were amended with GAC, and the potential of the working electrode was alternated between +0.4 and -0.4 V vs. SHE using a potentiostat. Six frequencies ranging from 0.5 to 12.0 days were evaluated and compared with two control conditions with constant potentials. The community structure and activity were analyzed by sequencing 16S rRNA gene and transcript amplicons, complemented by Bayesian network analysis to explore potential causal relationships among community members and environmental drivers. The findings of this study are expected to provide mechanistic insights into the enrichment of electrotrophic methanogens and electroactive partners, based on which the strategy for achieving robust, high-rate electro-methanogenesis can be optimized.

## 2. Materials and Methods

### 2.1 Reactor setup and operation

H-type bioelectrochemical systems were constructed based on a previous study (Supporting Information (SI) Method and Figure S1).^33^ The working-electrode chamber was inoculated with 10% (v/v) anaerobic digester sludge supernatant obtained from a local wastewater treatment plant and supplemented with synthetic soft drink wastewater containing 3,000 mg/L fructose, 2,200 mg/L polyethylene glycol 200, and 5 mM sodium bicarbonates as carbon sources, buffered with 0.1 M phosphate buffer saline.^16, 39^ The reactors were operated in a continuous mode with a hydraulic retention time (HRT) of 3.5 days. The working electrode potential was alternated between +0.4 and -0.4 vs. SHE to avoid abiotic oxygen and hydrogen evolution, respectively. Six alternating-polarity frequencies (0.5, 1, 2, 4, 6, 12 d) were tested along with two control conditions (constant positive control +0.4 V vs. SHE, and constant negative control -0.4 V vs. SHE). The experiments contained one start-up stage and three enrichments. Each stage lasted for 24.5 days (seven cycles), after which 2.5 g (5% w/w) of the cultured GAC was transferred into a new reactor containing 47.5 g fresh GAC to start a new enrichment stage.

### 2.2 Chemical and electrochemical analyses

Biogas was collected at the end of each HRT (3.5 days) using a gas-tight syringe and normalized to substrate volume and GAC mass (L/L/kg). The methane content was quantified using gas chromatography-mass spectrometry (8860/5977B, Agilent Technologies) equipped with an Agilent CP-Molsieve 5A column (25 m × 0.25 mm). Effluent samples were collected concurrently in gas-tight bags for determination of soluble chemical oxygen demand (sCOD) using a HACH colorimetric digestion kit (HACH DR6000) and pH using a Mettler-Toledo SevenCompact™ pH meter. Electricity generation because of applied potential was continuously monitored by the potentiostat software (VSP-300, Biologic). Methane yield was calculated as described previously:^46^

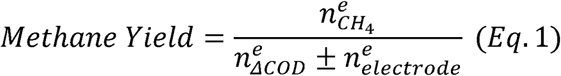

where 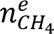, 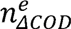, and 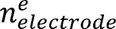 the number of electrons transferred for methane formation, the number of electrons derived from soluble COD removal, and the number of electrons transferred to/from the electrode, respectively:

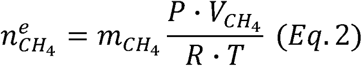

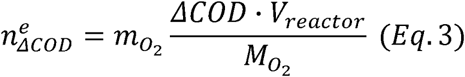

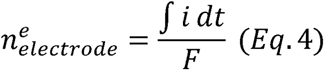

where m_CH4_ (8) is the mole of electron transferred per mole of CH_4_ formed, P (atm) is the measurement pressure, V_CH4_ is the volume of methane collected over each cycle (3.5 days), R (0.08206 L·atm/mol/K) is the ideal gas constant, T (K) is the measurement temperature, m_O2_ (4) is the mole of electron transferred per mole of oxygen consumed, ΔCOD (g/L) is the COD removal, V_reactor_ (0.085 L) is the reactor working volume, M_O2_ (32 g/mol) is the molecular weight of oxygen, i (A) is the electricity generated under the applied potential, and F (96485.3 C/mol) is the Faraday constant. Under the cathodic potential, additional electrons were supplied from the electrode to methanogenesis, and therefore, the number of electrons transferred was added to the theoretical number of electrons available from COD removal, as shown in Eq. 1. Under the anodic potential, part of the electrons released from substrate oxidation were transferred to the electrode and were therefore subtracted. Temporal reactor-performance data were analyzed using generalized least-squares models with a first-order autoregressive correlation structure [GLS-AR(1)] to account for serial dependence among repeated measurements from the same reactor. Planned contrasts were used to compare predefined reactor groups, and statistical significance was assessed at *p* < 0.05.^47^

### 2.3 Biomass sampling and amplicon sequencing

During each enrichment, 4 g of GAC was collected at the end of each HRT (3.5 days) from the working-electrode chamber and divided into four aliquots (two for DNA and two for RNA extraction). Samples for RNA extraction were pretreated with RNAlater™ (Thermo Fisher Scientific) to prevent degradation, and all samples were stored at -80 °C until nucleic acid extraction. Genomic DNA was extracted using the MPbio FastDNA Spin Kit (MP Biomedicals), and total RNA was isolated using the QIAGEN RNeasy PowerSoil Total RNA Kit (Qiagen). Residual DNA was removed using the QIAGEN DNA-free Kit, followed by reverse transcription with the SuperScript IV VILO Master Mix (Thermo Fisher Scientific) to obtain complementary DNA (cDNA). The quality and concentration of nucleic acids were verified using a NanoDrop ND-2000 spectrophotometer and Qubit™ dsDNA Quantification Assay (Thermo Fisher Scientific). The V4 region of the 16S rRNA gene and rRNA was amplified by polymerase chain reaction (PCR) using the primer pair 515F (GTGCCAGCMGCCGCGGGTAA) and 806R (GGACTACHVGGGTWTCTAAT), with sample-specific barcodes.^48^ The PCR products were subjected to quality assessment by gel electrophoresis, and amplicons with an expected length of approximately 300 bp were purified using ChargeSwitch Nucleic Acid Purification Technology (Invitrogen) and pooled in equimolar concentrations based on fluorescence-based quantification using a NanoDrop ND-3300 fluorospectrometer (Thermo Fisher Scientific). The pooled library was subsequently quantified with a KAPA Illumina Library Quantification Kit (Kapa Biosystems) and sequenced on an Illumina MiSeq platform. Paired-end reads were merged and denoised in QIIME 2, and amplicon sequence variants (ASVs) were generated using DADA2.^49, 50^ Taxonomic assignment was performed using the QIIME 2 feature-classifier plugin against the SILVA database, with the classifier trained on 95% reference sequences.^51^

### 2.4 Microbial community analysis

After removal of singleton reads, statistical analyses were conducted in R software (v4.3.1) using the vegan and phyloseq packages. Principal coordinate analysis (PCoA) based on Bray-Curtis dissimilarities were applied to assess microbial diversity and clustering. The significance of community separation was verified by permutational multivariate analysis of variance (PERMANOVA) with 999 permutations (*p* < 0.05). Core operational taxonomic units (OTUs) were identified based on average relative abundance and occurrence across all samples.^32^ A phylogenetic tree of the selected core OTUs was constructed using ARB software employing the neighbor-joining algorithm.^52^ Relative growth activity of a population was calculated by normalizing its 16S rRNA transcript abundance measured under the anodic and cathodic potentials against its corresponding 16S rRNA transcript abundance measured in the positive (+0.4 V vs. SHE) and negative (−0.4 V vs. SHE) controls, respectively. Redundancy analysis (RDA) was further performed in R using the relative activity of core populations derived from 16S rRNA transcript data and environmental variables. Before ordination, transcript data were subjected to Hellinger transformation, while environmental variables were standardized by z-score normalization.

### 2.5 Network analysis

Bayesian networks were constructed using the R package “bnlearn” to infer the interactions of the microbial populations.^53^ Relative abundances of the core OTUs derived from 16S rRNA gene sequencing were combined with three key environmental variables (methane percentage, cathodic current, and pH) into a single dataset. All variables were normalized to a 0-1 scale:

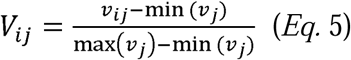

where V_ij_represents the normalized value of variable j in sample i, ν_ij_ is the observed value, and min (ν_j_)and max (ν_j_)are the minimum and maximum values of variable j, respectively. The network structure was learned using a hill-climbing algorithm, and model parameters were estimated by maximum likelihood.^54^ Model performance was evaluated using leave-one-out cross-validation.^55^ In each cross-validation cycle, the relative root mean square error (RMSE) between observed and predicted values was calculated:^56^

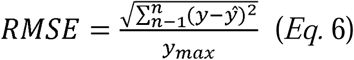

where y is the experimental value, ŷ is the predicted value, and y_max_ is the maximum experimental value. Null model predictions were generated by fixing the values of selected nodes (e.g., overall microbial abundance or environmental variables) to their mean values across all samples. Extended local similarity analysis (eLSA) were performed to identify pairwise associations among active microbial populations and environmental variables based on time-series data.^57^ The analysis used log-transformed 16S rRNA transcript abundance data and corresponding environmental parameters for each reactor. Pairwise associations were quantified using the local similarity (LS) score, and statistical significance was assessed by permutation testing followed by false discovery rate correction. Only associations with l LSl>0. 4 and *p* < 0.05 were retained for network construction and visualization.

## 3. Results and Discussion

### 3.1 Electro-methanogenic bioreactor performance

As a primary indicator of the performance of electro-methanogenic communities, cumulative biogas production (normalized to substrate volume and GAC mass) increased and stabilized as the enrichment proceeded (Figure 1A). After start-up, the cumulative biogas did not vary noticeably across alternation frequencies, ranging from 86 to 142 L/L/kg. As expected, the -0.4 V control produced the most biogas of 212 L/L/kg, and the +0.4 V control produced the least biogas of 30 L/L/kg. By the end of the first enrichment, 4-d alternation stood out with the highest cumulative biogas of 255 L/L/kg, which was 46 L/L/kg higher than that of the -0.4 V control. The 2- and 6-d alternation conditions produced similar cumulative biogas of ∼180 L/L/kg, while higher frequencies (i.e., 0.5- and 1-d) suppressed biogas production to ∼110 L/L/kg. The same trend was observed in the latter two enrichments, after which the 4-d group still achieved the highest final cumulative biogas production of ∼250 L/L/kg, followed by the -0.4 V control that produced ∼220 L/L/kg. The moderate conditions of 2- and 6-d alternation showed slightly enhanced performance with ∼195 L/L/kg, whereas the extreme conditions of 0.5-, 1-, and 12-d alternation produced significantly less biogas (< 170 L/L/kg) (GLS+AR (1) test, *p* < 0.05). The +0.4 V control always produced an order of magnitude less biogas (< 15 L/L/kg).

**Figure 1.**
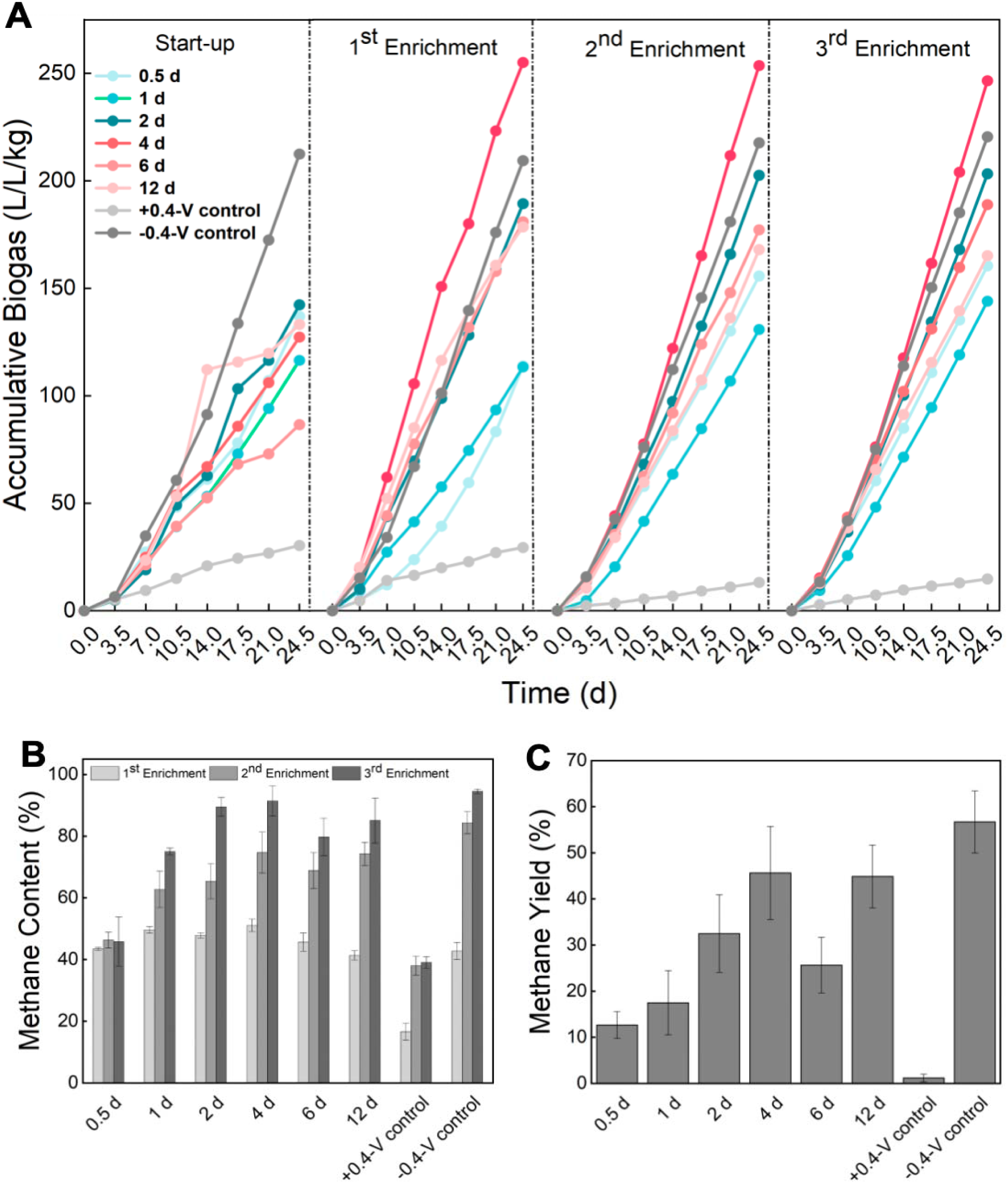
(A) Cumulative biogas production of six alternating frequencies and two control conditions in the start-up and three enrichments. (B) Methane content of the eight conditions over the three enrichments. (C) Methane yield of the eight conditions after the third enrichment.

Methane content as another important indicator showed a trend consistent with cumulative biogas production (Figure 1B). During the first enrichment, methane content remained moderate across all conditions, with the highest content of 52% achieved by 4-d alternation and the content under other conditions ranging from 43% to 49%. The -0.4 V control yielded a similar methane content of 46%, while the +0.4 V control had the lowest methane content of 18%. In the second enrichment, the methane content of the 4-d condition -0.4 V control increased substantially to 81% and 88%, respectively, followed by the 2- and 6-d conditions which achieved ∼75%. While the methane content under the 1- and 12-d conditions increased similarly to up to 75%, that of the 0.5-d alternation remained low at 49%. By the end of the third enrichment, the methane content of the 4-d condition and -0.4 V control further increased to 96% and 94%, respectively. Moderate conditions (2-, 6- and 12-d alternation) also improved methane content to up to 93%, while high frequencies (0.5- and 1-d alternation) significantly suppressed the performance (52% and 74%, respectively) (GLS+AR (1) test, *p* < 0.05). The +0.4 V control consistently show the lowest methane content of only 40%.

Regardless of the applied alternation frequencies, the cathodic current were consistently lower in magnitude than the anodic current (SI Figure S2). The mean cathodic current was observed to be 0.10 - 0.15 mA under higher frequencies (i.e., 0.5, 1, 2 and 4 d) but as low as 0.04 mA under lower frequencies (i.e., 6 and 12 d). Similarly low cathodic currents have been reported previously and may reflect the limited abundance and/or activity of cathode-associated electrotrophic methanogens.^58^ Consistent with this trend, mean anodic current was high under higher frequencies, particularly the 1-d condition (0.52 mA), followed by the 0.5- and 2-d conditions (0.34 mA). As the frequency was increased from 4 d to 12 d, the mean anodic current decreased from 0.21 mA to 0.10 mA. Anodic and cathodic currents together reflected the microbial activity, which subsequently resulted in variation in pH.^59, 60^ The lowest effluent pH of 6.3 was observed under the 4-d condition and the -0.4 V control (SI Figure S3). The other frequencies maintained similar pH values of approximately 6.6, whereas the +0.4 V control exhibited the highest mean pH of 7.2. Such acidification might result from volatile fatty acids production and/or proton release from anode-associated organic oxidation,^59, 61^ implying high activity of fermentative and electroactive populations.

This interpretation was supported by COD removal (SI Figure S3). The -0.4 V control achieved the highest and most stable COD removal of 52% across the three enrichments, and the 2- and 4-d conditions achieved similarly high COD removal of up to 50%. Consistent with the trends of other indicators, low frequencies (6 and 12 d) showed slightly lower COD removal of 40% by the end of the third enrichment, whereas high frequencies (0.5 and 1 d) suppressed COD removal to as low as 31%. The +0.4 V control consistently exhibited the lowest COD removal of 17%. Derived from methane content, current generation, and COD removal (Eq. 1), methane yield as a final indicator of the electro-methanogenic performance was again the highest in the -0.4 V control (Figure 1C), reaching on average 57%. Among the alternating polarity reactors, the 4- and 12-d conditions showed similar methane yields of 45%, which were significantly higher than the 33% under the 2- and 6-d conditions (GLS+AR (1) test, *p* < 0.05). In contrast, high frequencies (0.5 and 1 d) exhibited much lower methane yields of under 18%, and the +0.4 V control showed the lowest methane yield of only 1%. Overall, these results collectively demonstrate that 4-d alternation was the optimal frequency to build robust electro-methanogenic communities, while the communities assembled under high frequencies were less effective in coupling organic matter degradation, extracellular electron transfer, and methane production.

### 3.2 Electro-methanogenic community dynamics

Bray-Curtis dissimilarity-based PCoA revealed significant separation of the electro-methanogenic communities assembled with different alternation frequencies (Figure 2, PERMANOVA, *p* < 0.05). By the end of the third enrichment, the optimal communities under 4-d alternation formed a relatively compact and distinguishable cluster toward positive PC1 and negative PC2. The communities in the -0.4 V control were observed on the left side of those under 4-d alternation. In comparison, the sub-optimal communities under 2-d alternation shifted further left to the origin of the PCoA graph, whereas the other sub-optimal communities under 6-d alternation appeared on far-left end. On the other hand, the communities assembled with more extreme frequencies were either far away from the optimal communities (i.e., 0.5- and 12-d alternation and the +0.4 V control) or widely dispersed (i.e., 1-d alternation). For example, three of the four communities assembled with 1-d alternation were structurally similar to those from 2- and 4-d alternation, but the remaining one was found distantly separated in the upper left corner of the PCoA graph.

**Figure 2.**
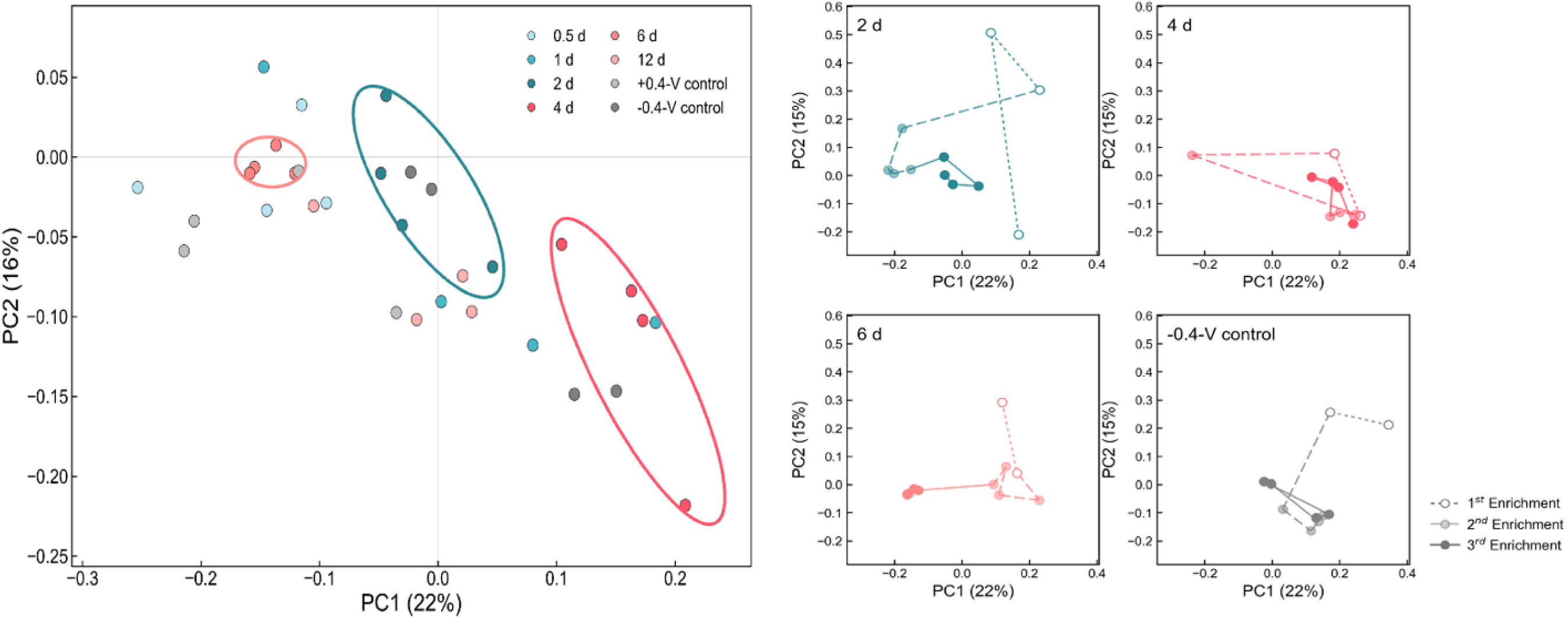
(A) Bray-Curtis dissimilarity-based PCoA of the electro-methanogenic communities assembled under 2-, 4- and 6-d alternation frequencies. For each frequency condition, samples were collected of the last four cycles of the third enrichment. (B) The temporal trajectories of community dynamics under 2-, 4- and 6-d alternation frequencies and -0.4v control in three enrichments.

The temporal trajectories across the three enrichments further revealed distinct dynamics of the communities assembled with different alternation frequencies (Figure 2 and SI Figure S4). The communities under 4-d alternation showed variation in strucutre in the early stage but quickly stabilized as the enrichment proceeded. Similarly, the -0.4 V control exhibited an early shift but became structurally stable in later enrichments. Other alternation frequencies, however, appeared to impose continuous impacts on community dynamics. For example, the communities under 2-d alternation shifted drastically from the bottom center of the PCoA graph (early first enrichment) over the upper right corner (late first enrichment) to the left center (second enrichment) and finally reached the origin (third enrichment). The communities under 6-d alternation were less dispersed but still noticeably differentiated across the three enrichments. Similar successional dynamics and enrichment-based differentiation were observed in communities under 0.5, 1- and 12-d alternation. Overall, the PCoA results suggest that 4-d alternation was the optimal frequency to stabilize electro-methanogenic community structure, while higher and lower frequencies might introduce strong disturbance to community assembly.

### 3.3 Electro-methanogenic community composition

Based on the criteria of average relative abundance greater than 0.75% and occurrence in more than 75% of the DNA samples, 24 OTUs were identified as core populations (Figure 3), collectively accounting for approximately 70% of the total relative abundance. A close inspection of these core populations revealed three major functional guilds: methanogens, putative electroactive bacteria, and fermentative and syntrophic bacteria.

**Figure 3.**
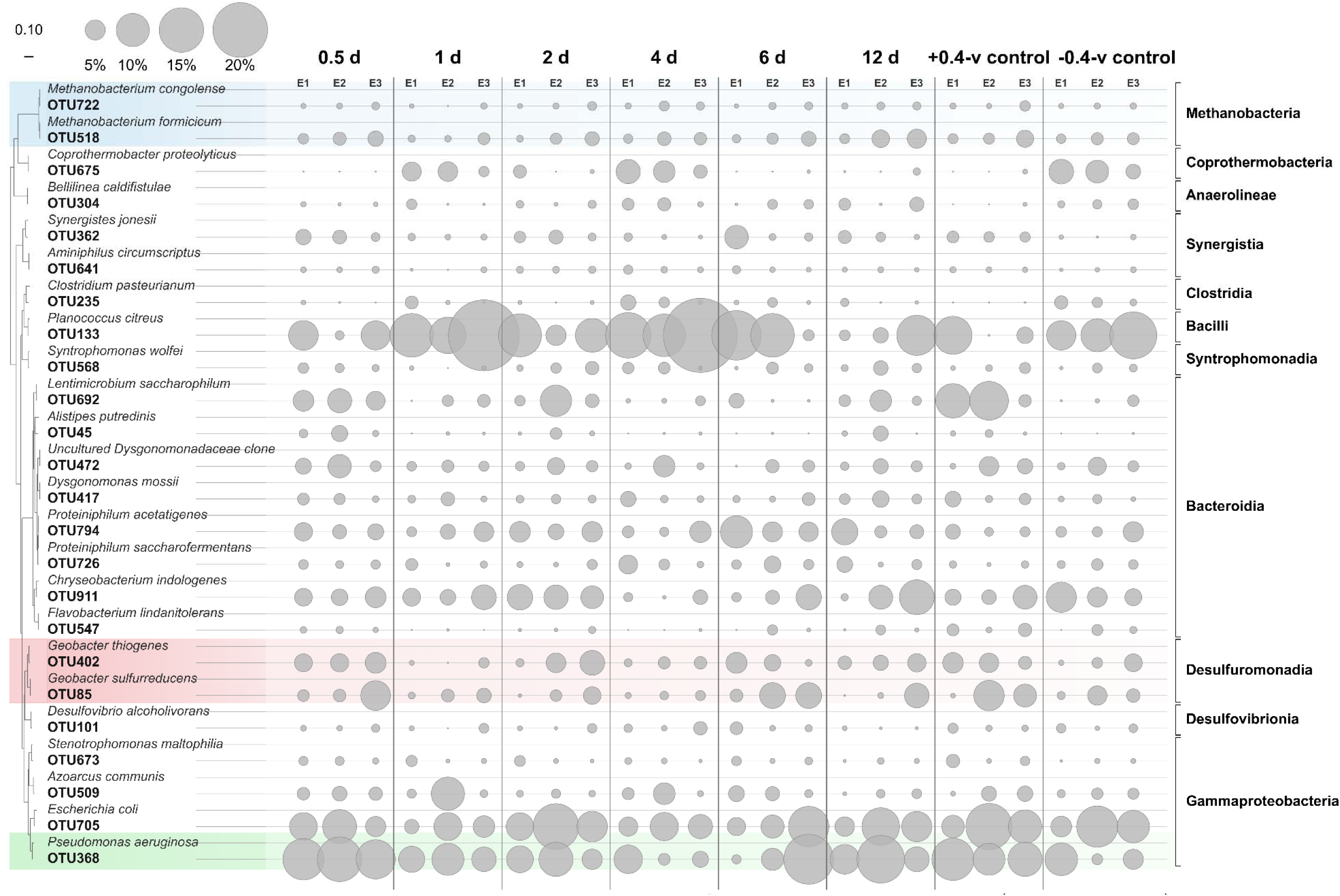
Phylogenetic tree and relative abundance of the 24 core OTUs in three enrichments (E1, E2, and E3). The abundance in each enrichment was obtained by averaging the abundances from the final three sampling cycles of each enrichment.

The methanogenic core populations were represented by OTU722 and OTU518, which were phylogenetically related to *Methanobacterium congolense* and *Methanobacterium formicicum*, respectively. The abundance of *M. congolense*-related OTU722 remained consistently low across different frequencies, reaching 2% under 4-d alternation in the later enrichments and approximately 1% under other conditions. *M. formicicum*-related OTU518 accounted for around 3% of the community under 4-d alternation and the -0.4 V control across the three enrichments. Under other conditions, OTU518 showed progressive enrichment, with the highest abundance of 5% observed under 12-d alternation in the third enrichment. The persistence of *Methanobacterium* populations under alternating polarity is consistent with previous observations of high abundances of *Methanobacterium* in the bioanode of bioelectrochemical systems and in the biocathode of methanogenic bioreactors.^19, 33, 39^ While *Methanobacterium spp.* are known to perform conventional H_2_- or formate-dependent methanogenesis, recent studies have proposed their ability of extracellular electron uptake from cathodes and electroactive partners.^29, 62–64^

Three putative electroactive bacteria were observed with distinct abundance under different frequencies and enrichments. Among them, *Geobacter thiogenes*-related OTU402 showed stable abundance of 1-3% under 4-d alternation across the three enrichments. The highest abundance of this population was observed under 2-d alternation (7% in the third enrichment), and other alternation conditions slightly favored its growth (2-5% in the third enrichment). It was interesting to observe the enrichment of OTU402 in the -0.4 V control and a declining trend in the +0.4 V control across enrichments. Similarly, *Geobacter sulfurreducens-*related OTU85 maintained a relatively stable abundance under 4-d alternation across the three enrichments. Its abundance in the third enrichment ranged from 3% under 1-d alternation to 9% under 0.5-d alternation. Both the -0.4 V and +0.4 V control appeared to favor its growth. The third electroactive candidate, *Pseudomonas aeruginosa*-related OTU368, had an average abundant of 5% under 4-d alternation. The highest abundance of this population was observed under 6-d alternation (18% in the third enrichment), and its abundance under other conditions, including the -0.4 V (5%) and +0.4 V control (12%), was comparably high. *Geobacter spp.* are well known for their electroactivity,^61^ and their interactions with electrotrophic methanogens have also been extensively reported.^65^ On the other hand, *Pseudomonas spp.* have been reported to mediate extracellular electron transfer through electron shuttles (e.g., phenazines),^66^ but their the dominance in electro-methanogenic communities is documented for the first time in this study.

Several fermentative and syntrophic populations were consistently observed in the core community. Among them, *Planococcus citreus*-related OTU133 was dominant with 10-18% of the community and reached as high as 30% under 4-d alternation. *Lentimicrobium saccharophilum*-related OTU692 was abundantly present with approximately 9% and 12% under the 2-d condition and the +0.4 V control, respectively. *Proteiniphilum acetatigenes-*related OTU794 showed relatively even abundances of 3-7% across the frequencies and enrichments. Based on previous studies on isolates,^67, 68^ these populations are speculated to perform hydrolysis and fermentation, thereby supplying metabolic intermediates to electroactive and methanogenic populations. Additionally, *Coprothermobacter proteoly*ticus-related OTU675, a potential thermophilic proteolytic fermenter capable of degrading proteins and peptides and producing H□ and acetate,^69^ exhibited up to 7% under 4-d alternation and in the -0.4 V control. OTU568, which was related to *Syntrophomonas wolfei* (a well-established syntrophic fatty-acid oxidizer that could provide H_2_ and formate to methanogenic partners)^70, 71^, was found with an abundance of < 3% throughout the enrichments. Similarly, OTU101 also showed an abundance of < 3% and was affiliated with *Desulfovibrio*, some of which could participate in H_2_- and formate-dependent syntrophic metabolism.^70, 72^

### 3.4 Electro-methanogenic population activity

The growth activity of the 24 core populations, represented by their relative 16S rRNA transcript abundances, showed clear frequency-dependent patterns (Figure 4A and SI Figure S5). For the two methanogens, 4-d alternation favored the activity of *M. congolense*-related OTU722 (3%) more than that of *M. formicicum*-related OTU518 (1%). Similarly, OTU722 was significantly more active than OTU518 under most of the conditions, particularly under 2-d alternation (9% vs. 1%, GLS+AR (1) test, *p* < 0.05). When comparing the two *Geobacter* populations, alternating polarity stimulated the activity of *G. sulfurreducens*-related OTU85 over *G. thiogenes*-related OTU402. For example, the activity OTU85 and OTU402 was 11% and 4%, respectively, under 4-d alternation (GLS+AR (1) test, *p* < 0.05). Consistent with this pattern, higher OTU85 activity was observed under the 2- and 6-d conditions and both constant-potential controls (8 - 28%), whereas the activity of OTU402 was comparable under the 2-d condition and the +0.4 V control (5%) but was much lower under the 6-d condition and the -0.4 V control (< 2%). *P. aeruginosa*-related OTU368 also showed higher activity under the 4-d condition and the -0.4 V control (14%) but much lower activity under the 2- and 6-d conditions (1%). *C proteoly*ticus-related OTU675 was concurrently active under 4-d alternation and the -0.4 V control (13%), while other fermentative and syntrophic populations remained relatively inactive.

**Figure 4.**
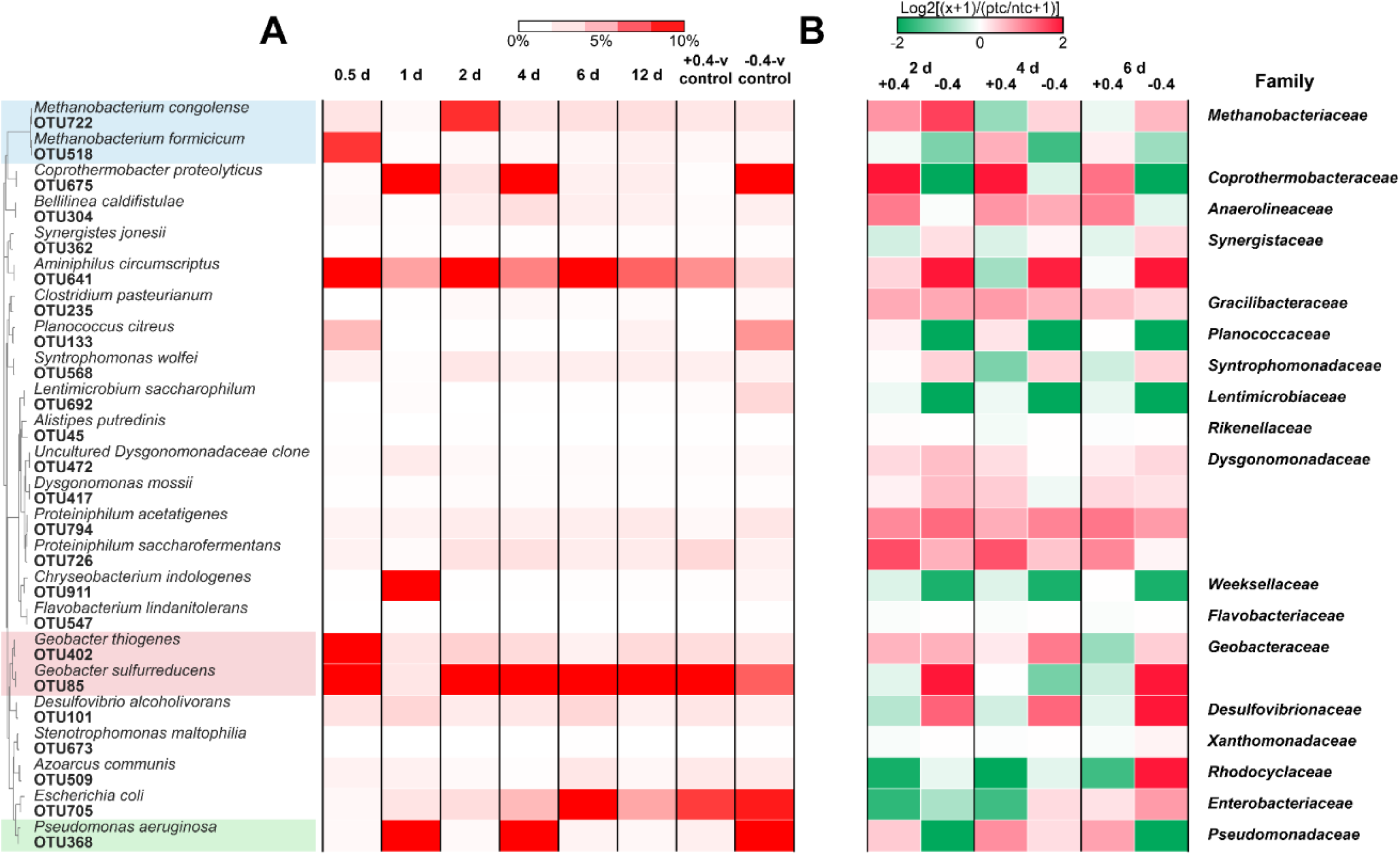
(A) Growth activity of the 24 core populations in the third enrichment, represented by mean relative 16S rRNA transcript abundance over the last four cycles of the third enrichment. (B) Activity regulation under 2-, 4- and 6-d frequency in the third enrichment, calculated as the ratio of the mean relative 16S rRNA transcript abundances during the positive- and negative-potential cycles normalized to those in the corresponding +0.4 V and -0.4 V controls, respectively.

The relative activity of the 24 core populations under 2-, 4- and 6-d alternation normalized to the controls further demonstrated distinct potential-dependent responses (Figure 4B). Compared with *M. congolense*-related OTU722 which was stimulated by -0.4 V (1.3 - 3.1 folds), *M. formicicum*-related OTU518 was consistently suppressed by -0.4 V (0.4 to 0.6-fold), indicating less reliance of OTU518 on the cathodic electrode. For the two *Geobacter*, *G. thiogenes*-related OTU402 was stimulated by -0.4 V with the strongest up-regulation observed under 4-d alternation (2.2 folds), whereas *G. sulfurreducens*-related OTU85 was suppressed by -0.4 V under 4-d alternation (0.5-fold). Interestingly, the activity of both *Geobacter* populations was not stimulated by anodic +0.4 V, which in principle served as a stable electron sink for electroactive bacteria, implying their weak reliance on the anodic electrode. In comparison, *P. aeruginosa*-related OTU368 was consistently stimulated by +0.4 V across all three frequencies (1.4 - 2.0 folds) but suppressed by -0.4 V, implying its electroactivity. Within the fermentative and syntrophic populations, *P. acetatigenes*-related OTU794 was broadly stimulated under both potentials across the tested frequencies. The activity of *C. proteolyticus*-related OTU675 showed consistent up-regulated by +0.4 V and down-regulated by -0.4 V, whereas that of *Desulfovibrio*-related OTU101 showed the opposite pattern across the three frequencies.

RDA based on relative 16S rRNA transcript revealed distinct associations among electrochemical conditions, population activity, and reactor performance (SI Figure S6). As expected, the cathodic and anodic currents extended in contrasting directions. The cathodic current showed a strong association with community transcriptional patterns (F = 1.7, *p* < 0.05), with *M. formicicum*-related OTU518 positioned along the positive direction. This methanogen was also positively associated with biogas production and methane production (F = 2.7, *p* < 0.05). The anodic current was positively associated with *G. sulfurreducens*-related OTU85, which was comparatively more oriented toward methane production. *G. thiogenes*-related OTU402, on the other hand, was separated from the methanogen OTU85 but was broadly corresponding to the pH gradient. The third putative electroactive bacteria, *P. aeruginosa*-related OTU368, showed a noticeable association with anodic current, confirming its electroactivity.

### 3.5 Electro-methanogenic population interactions

Bayesian networks constructed using the relative 16S rRNA gene abundances of the core OTUs together with key environmental variables revealed noticeable interactions between core populations and electrochemical conditions (Figure 5A, SI Table S1). The network inferred the connection between *M. congolense*-related OTU722 and three parent nodes: current (coefficient 0.25), OTU641 (0.18), and OTU692 (−0.15). *M. formicicum*-related OTU518 was inferred to be associated with six parent nodes, including the positive associations with the methanogen OTU722 (0.77) and the two *Geobacter* OTU402 (0.35) and OTU85 (0.33). Moreover, OTU518 had a positive impact on methane production (0.34). As OTU518 connected other populations and methane production, predictive inference was performed on this population using leave-one-out cross-validation, and moderate predictive power for OTU518 abundance was achieved with the observed values of all of its parents (R² = 0.62 and RMSE = 12.4 ± 8.2%) (Figure 5B). Null predictions by fixing the abundance of *Geobacter* OTU85 (one of OTU518’s parent nodes) at zero while retaining the observed values of the remaining parent nodes only slightly decreased R² to 0.56 and increased RMSE to 13.4 ± 9.1% (Figure 5C). In contrast, fixing the abundance of *Geobacter* OTU402 at zero resulted in a significantly lower R² of 0.53 and higher RMSE of 13.9 ± 9.6% (*p* < 0.05, Figure 5D). A more pronounced deterioration of the prediction of OTU518 abundance was observed when COD was fixed at zero, with R² decreasing to 0.52 and RMSE increasing to 14.0 ± 8.2% (*p* < 0.05, Figure 5E). These null-prediction results implied the interactions between *Methanobacterium* OTU518 and *Geobacter* OTU402 during electro-methanogenesis.

**Figure 5.**
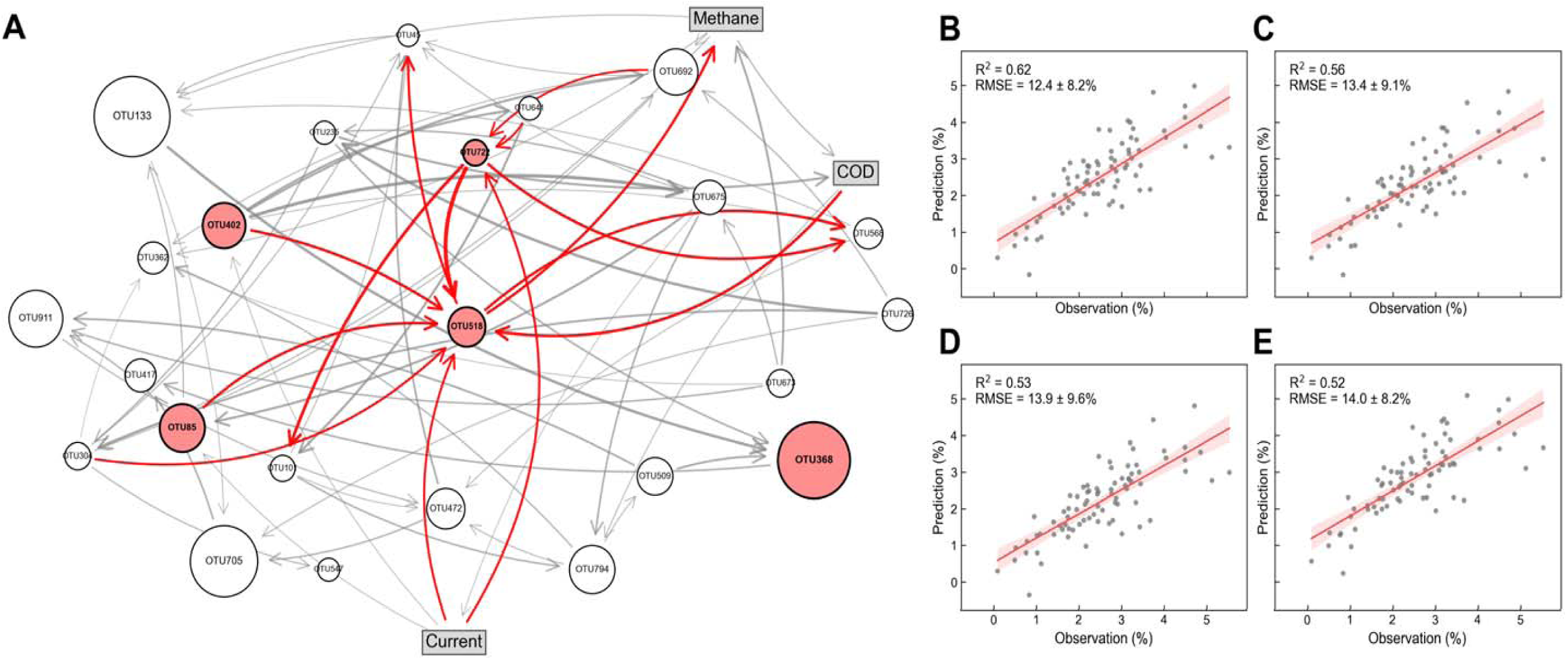
(A) Bayesian network constructed with the core populations (circle nodes) and three environmental factors (rectangular nodes). Prediction of the abundance of *Methanobacterium* OTU518 was performed with (B) the observed values of all its parent nodes, or after fixing (C) OTU85, (D) OTU402, and (E) COD at zero while retaining the observed values of all other parent nodes.

The integrated eLSA network further revealed the contribution of electrochemical conditions and microbial interactions to methane production (SI Figure S7). The cathodic current was located at the network periphery and was directly associated only with OTU45, through which it was indirectly connected to the rest of the microbial network. *M. congolense*-related OTU722 showed a direct positive association with methane production (LS = 0.45) and a negative association with *P. citreus*-related OTU133 (LS = -0.48). *M. formicicum*-related OTU518 was indirectly connected to the electrochemical conditions through neighboring populations and was indirectly linked to methane through the broader microbial network. The two *Geobacter* populations displayed distinct network connectivity. *G. sulfurreducens*-related OTU85 was positively associated with methane production (LS = 0.54), *Desulfovibrio*-related OTU101 (LS = 0.40) and *M. congolense*-related OTU722 (LS = 0.58). On the other hand, *G. thiogenes*-related OTU402 showed a negative association with OTU85 (LS = -0.41) and connected through intermediate populations to the OTU518-associated network. *P. aeruginosa*-related OTU368 was negatively associated with pH (LS = -0.53), linking its activity to reactor conditions in addition to its association with OTU518. Among the fermentative and syntrophic populations identified above, *C. proteolyticus*-related OTU675 was directly associated with COD (positive, LS = 0.41), whereas *L. saccharophilum*-related OTU692 was directly associated with anodic current (positive/negative, LS = 0.40).

### 3.6 Perspectives

Electro-methanogenesis as a new pathway to enhance methane recovery has attracted increasing interest. Conventional strategies to promote electro-methanogenesis, including cathodic enrichment, microbial electrolysis, and conductive-material amendment, are not efficient due to their inability to simultaneously enrich the functional populations involved in electro-methanogenesis.^23, 28, 73^ Alternating polarity was recently developed as a community-engineering strategy in which periodic anodic and cathodic stimulation could provide selective conditions for both electroactive bacteria and electrotrophic methanogens.^33^ During alternating polarity, the assembly of electro-methanogenic communities depends not only on the applied potentials but also on the frequency of the alternation. Excessively frequent potential changes can disrupt electroactive biofilms and/or cannot provide sufficient time for microbial adaptation,^74^ whereas prolonged polarization may favor one functional guild at the expense of others.^33^ ^75^ The present study systematically evaluated a wide range of alternation frequencies and confirmed that high frequencies of 0.5-1 d substantially lowered biogas production and methane content. An alternation frequency of 4 d was found to be the optimal frequency for building robust electro-methanogenic communities. Compared to our previous study, the optimized frequency increased biogas production by over 100% and achieved approximately 96% methane content with 45% methane yield.

Alternation frequency strongly influenced the structure of electro-methanogenic communities and the activity of functional populations. The dominance and high activity of putative fermentative populations *P. citreus*-related OTU133 and *C. proteolyticus*-related OTU675 under 4-d alternation suggest faster upstream organic-matter conversion.^76^ The broad stimulation of *P. acetatigenes*-related OTU794 under both potentials further implies sustained fermentative substrate conversion during polarity alternation (Figures 3 and 4).^67, 75^ *P. aeruginosa*-related OTU368 was consistently stimulated by the anodic condition potentially through phenazine-mediated extracellular electron transfer (Figure 4B; SI Figures S4 and S6),^66, 77^ but its involved in methanogenesis warrant further investigation. In contrast, the two *Geobacter* populations showed relationships more consistent with contributions to methanogenesis: *G. sulfurreducens*-related OTU85 was associated with anodic current and methane production in the RDA and connected to methane, methanogenic OTU722, and syntrophic OTU101 in the eLSA network, whereas *G. thiogenes*-related OTU402 was cathodically stimulated under 4-d alternation and contributed substantially to Bayesian prediction of methanogenic OTU518 (Figures 4 and 5; SI Figures S4 and S6). The two methanogens were both from the genus *Methanobacterium* and potentially carried out both electrotrophic and hydrogenotrophic methanogens.^78–80^ The high activity, cathodic stimulation, and associations with methane and OTU85 imply the ability of OTU722 to receive reducing equivalents from the cathode or electroactive partners. OTU518, on the other hand, was anodically stimulated but associated with cathodic current, biogas production, both *Geobacter* populations, and OTU722, suggesting that its activity may be influenced by community-mediated interactions than by direct cathodic electron uptake (Figures 4 and 5; SI Figures S4 and S6).

While previous work has shown that the magnitude of anodic and cathodic potentials can shape electroactive community,^33^ the present study has demonstrated the impacts of duration of each potential on electro-methanogenic performance, and community structure, and microbial activity and interaction. Based on these results, future studies can focus on the simultaneous optimization of applied potential and alternation frequency. The interplay between these two types of operating conditions will collectively affect the abundance and activity of fermentative, electroactive, methanogenic populations. For example, asymmetric anodic and cathodic potentials can be applied with asymmetric frequencies (e.g., +0.8 V vs. SHE for two days and -0.4 V vs. SHE for six days) by considering the growth rates of electroactive bacteria and electrotrophic methanogens. Beyond mechanistic optimization, the practical application of alternating polarity will depend on whether the additional methane recovery compensates for the electrical energy required to establish and maintain the community. Reactor configuration, energy requirements, and scale-up remain major challenges for practical electro-methanogenesis.^69, 81^ Long-term experiments using real wastewater or anaerobic-digestion feedstocks are therefore needed to evaluate community stability under fluctuating conditions and to determine the necessity of electrical stimulation after temporary operational interruption.

## 4. Conclusions

In this study, the impact of alternating polarity frequencies on the assembly of electro-methanogenic communities was studied. A mild frequency of 4 d consistently led to the highest biogas production and methane content over three successive enrichments. DNA/RNA-based analyses showed that the optimal polarity alternation frequency promoted the co-enrichment and sustained the activity of putative electroactive bacteria and electrotrophic methanogens, enabling stable DIET-driven syntrophic interactions. Bayesian and local similarity network analysis further confirmed that methanogen dynamics were jointly regulated by electrochemical conditions and specific electroactive partners, emphasizing syntrophy-mediated electron transfer as the dominant pathway. Overall, these results provide mechanistic guidance for tuning alternating polarity strategies to enhance the stability, efficiency, and methane quality of electro-methanogenic systems.

## Acknowledgment

This work was supported by the National Science Foundation (CBET-2128365) and the Global Energy Ecosystems (GE^2^) Research Development Program of the Office of Research, Innovation & Economic Development at the University of Tennessee, Knoxville.

