## Supplementary Information for "Effects of Alternating Polarity Frequency on the Assembly of Electro-Methanogenic Microbial Communities"

**Frequency-Modulated Alternating Potentials Promote Methanogenesis in Anaerobic Systems**

Type of contribution***: Research Article***

^*^ Corresponding authors.

Heyang Yuan:

Qiang He:

Table S1. Directed edge coefficients among core microbial OTUs and environmental variables in the final Bayesian network.

|  | **Parent node** | **Child nodes** | **Coefficient** | **Direction** |
| --- | --- | --- | --- | --- |
| 1 | Methane | COD | 0.303 | Positive |
| 2 | OTU235 | COD | 0.528 | Positive |
| 3 | OTU675 | Current | -0.177 | Negative |
| 4 | OTU518 | Methane | 0.342 | Positive |
| 5 | OTU673 | Methane | -0.520 | Negative |
| 6 | OTU641 | OTU101 | 0.559 | Positive |
| 7 | OTU722 | OTU101 | 0.475 | Positive |
| 8 | Methane | OTU133 | 0.245 | Positive |
| 9 | OTU45 | OTU133 | -0.284 | Negative |
| 10 | OTU568 | OTU133 | -0.236 | Negative |
| 11 | OTU85 | OTU133 | -0.293 | Negative |
| 12 | OTU675 | OTU235 | 0.397 | Positive |
| 13 | OTU726 | OTU235 | 0.646 | Positive |
| 14 | Methane | OTU304 | 0.278 | Positive |
| 15 | OTU235 | OTU304 | 0.352 | Positive |
| 16 | OTU726 | OTU304 | 0.504 | Positive |
| 17 | Methane | OTU362 | -0.249 | Negative |
| 18 | OTU304 | OTU362 | -0.149 | Negative |
| 19 | OTU673 | OTU362 | 0.191 | Positive |
| 20 | OTU692 | OTU362 | 0.245 | Positive |
| 21 | OTU794 | OTU362 | 0.335 | Positive |
| 22 | OTU133 | OTU368 | -0.649 | Negative |
| 23 | OTU235 | OTU368 | -0.426 | Negative |
| 24 | OTU509 | OTU368 | -0.490 | Negative |
| 25 | Current | OTU402 | 0.199 | Positive |
| 26 | OTU368 | OTU417 | 0.422 | Positive |
| 27 | OTU705 | OTU417 | 0.429 | Positive |
| 28 | OTU101 | OTU45 | -0.236 | Negative |
| 29 | OTU304 | OTU45 | -0.274 | Negative |
| 30 | OTU472 | OTU45 | 0.473 | Positive |
| 31 | OTU518 | OTU45 | 0.308 | Positive |
| 32 | OTU675 | OTU45 | -0.254 | Negative |
| 33 | OTU692 | OTU45 | 0.237 | Positive |
| 34 | OTU101 | OTU472 | -0.217 | Negative |
| 35 | OTU568 | OTU472 | 0.226 | Positive |
| 36 | OTU794 | OTU472 | -0.247 | Negative |
| 37 | OTU911 | OTU472 | -0.290 | Negative |
| 38 | OTU794 | OTU509 | -0.233 | Negative |
| 39 | COD | OTU518 | -0.387 | Negative |
| 40 | Current | OTU518 | 0.228 | Positive |
| 41 | OTU304 | OTU518 | 0.229 | Positive |
| 42 | OTU402 | OTU518 | 0.350 | Positive |
| 43 | OTU722 | OTU518 | 0.769 | Positive |
| 44 | OTU85 | OTU518 | 0.332 | Positive |
| 45 | OTU304 | OTU547 | -0.290 | Negative |
| 46 | OTU402 | OTU568 | 0.220 | Positive |
| 47 | OTU518 | OTU568 | 0.326 | Positive |
| 48 | OTU722 | OTU568 | 0.366 | Positive |
| 49 | OTU402 | OTU641 | 0.546 | Positive |
| 50 | OTU402 | OTU675 | -0.757 | Negative |
| 51 | OTU673 | OTU675 | -0.350 | Negative |
| 52 | Methane | OTU692 | -0.227 | Negative |
| 53 | OTU304 | OTU692 | -0.275 | Negative |
| 54 | OTU402 | OTU692 | 0.551 | Positive |
| 55 | OTU726 | OTU692 | -0.322 | Negative |
| 56 | OTU133 | OTU705 | -0.200 | Negative |
| 57 | OTU472 | OTU705 | 0.361 | Positive |
| 58 | OTU726 | OTU705 | -0.256 | Negative |
| 59 | Current | OTU722 | 0.250 | Positive |
| 60 | OTU641 | OTU722 | 0.181 | Positive |
| 61 | OTU692 | OTU722 | -0.145 | Negative |
| 62 | COD | OTU794 | 0.232 | Positive |
| 63 | OTU101 | OTU794 | 0.398 | Positive |
| 64 | OTU675 | OTU794 | -0.430 | Negative |
| 65 | Current | OTU85 | 0.201 | Positive |
| 66 | OTU675 | OTU85 | -0.536 | Negative |
| 67 | OTU509 | OTU911 | -0.467 | Negative |
| 68 | OTU673 | OTU911 | 0.390 | Positive |
| 69 | OTU85 | OTU911 | 0.360 | Positive |

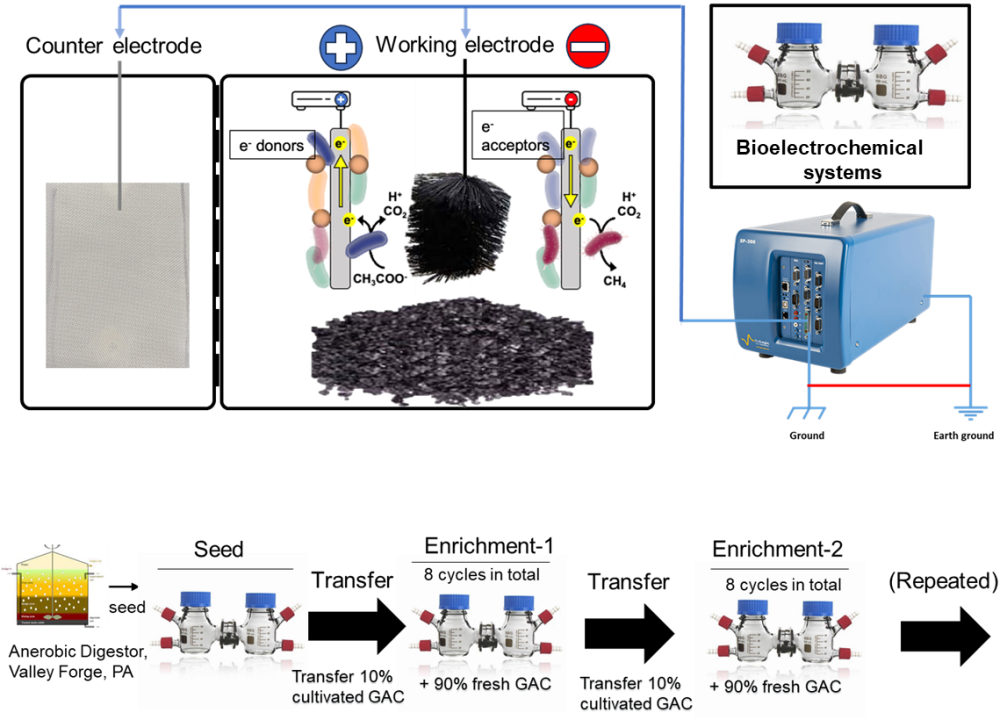

Figure S1. Schematics of the bioelectrochemical systems and cultivation procedure. H-type bioelectrochemical systems were constructed using two identical glass bottles (120 mL each) separated by a proton exchange membrane. In the working-electrode chamber, a stainless-steel cage was packed with 50 g of GAC to function as the working electrode, resulting in an 85 mL liquid volume. An Ag/AgCl_2_ reference electrode (+0.197V vs. SHE) was installed on the top cap without direct contact with the working electrode. A gas bag equipped with a septum was also installed on the top cap to allow biogas and effluent collection without air intrusion. The working-electrode chamber was maintained under anaerobic conditions (N₂/CO₂, 80:20 v/v) and at 35 ± 1 °C by wrapping thermostatically controlled mats around glass bottle. The solution in the working-electrode chamber was recirculated using a peristaltic pump (Masterflex L/S, Cole-Parmer) at 46 mL/min to ensure mixing. The counter-electrode chamber contained a stainless-steel mesh and 0.1 M phosphate buffer saline.

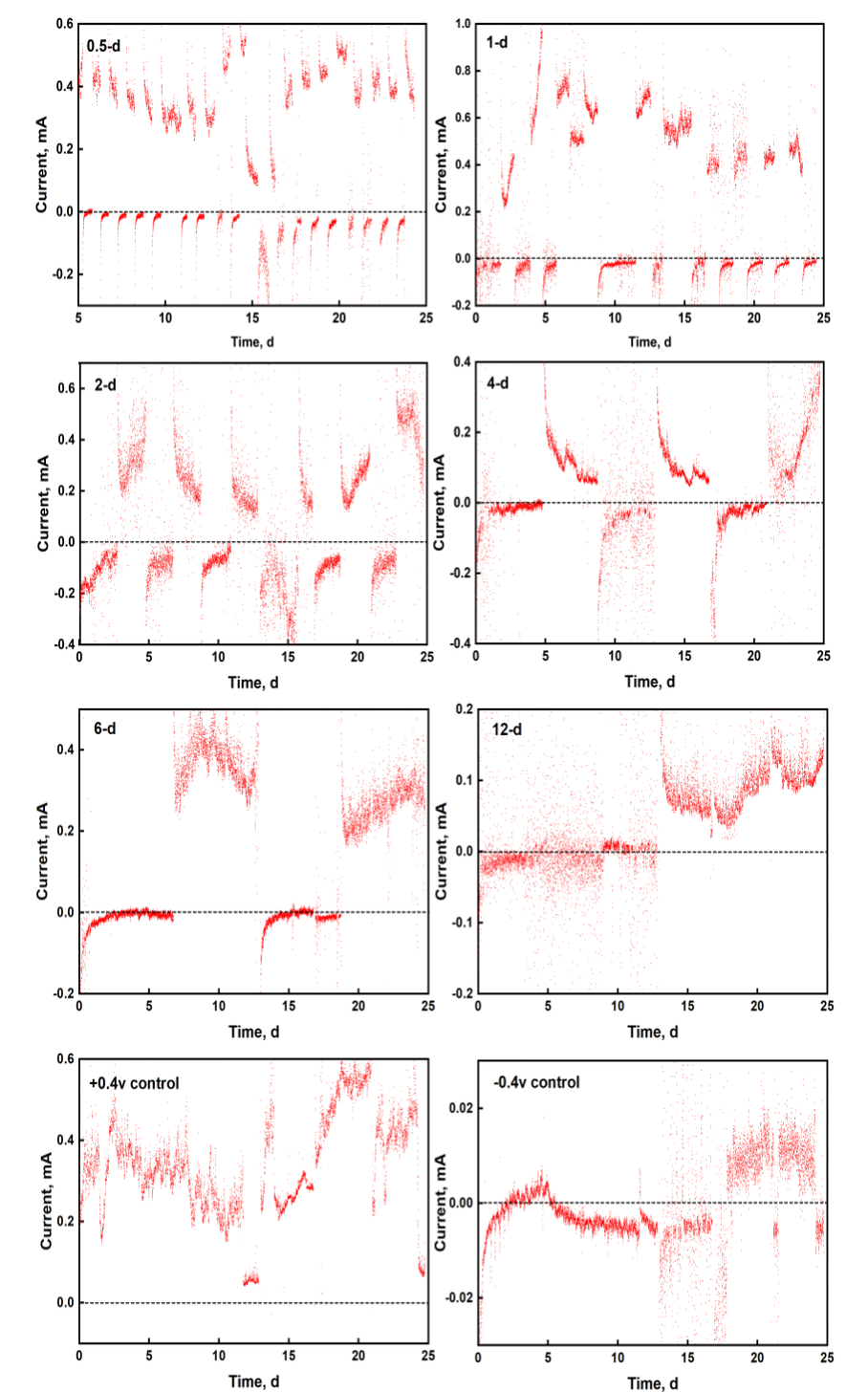

Figure S2. Voltametric responses of six alternating conditions and two controls in the 3rd enrichment.

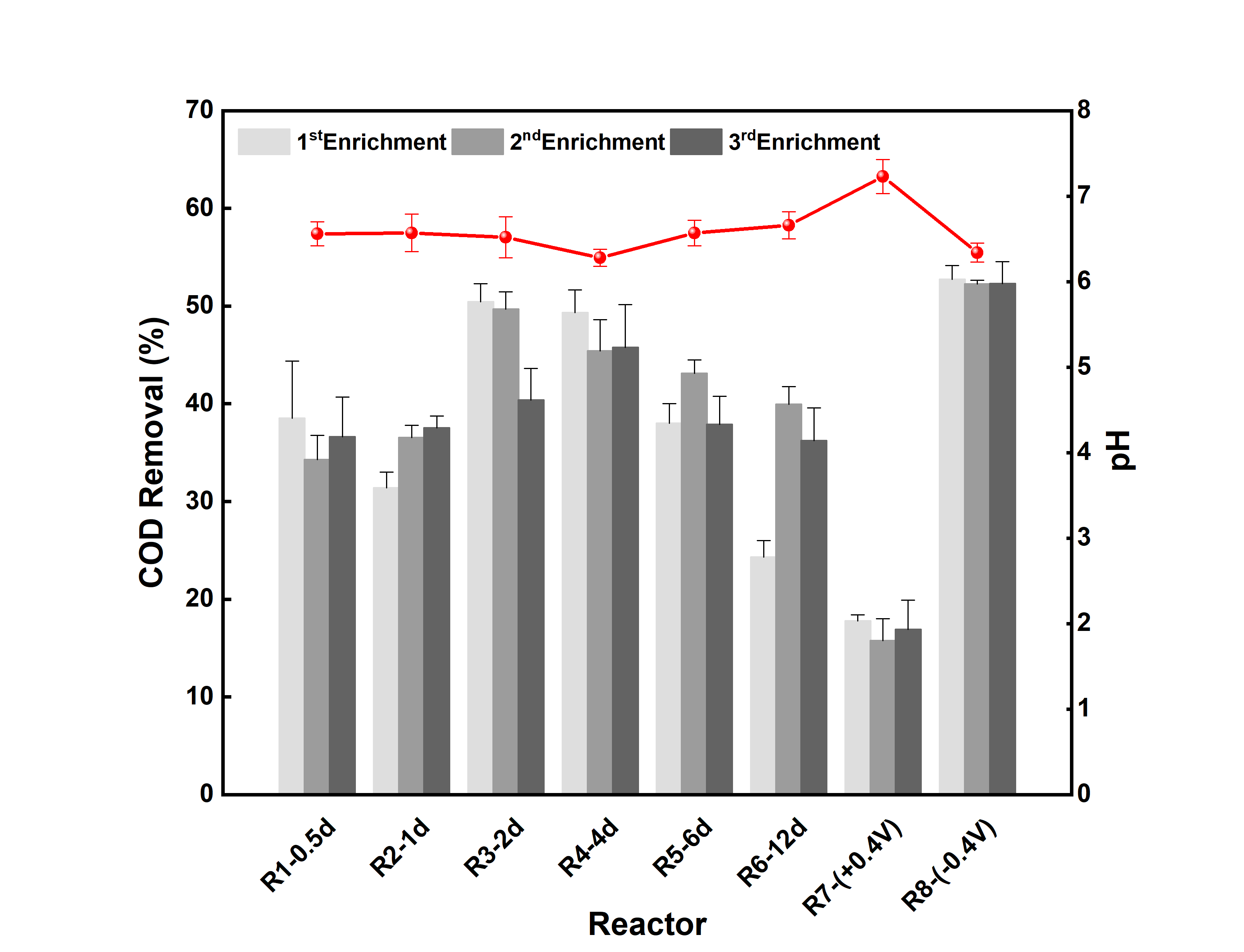

Figure S3. Soluble COD removal of eight conditions in the three enrichment and pH in the 3^rd^ enrichment.

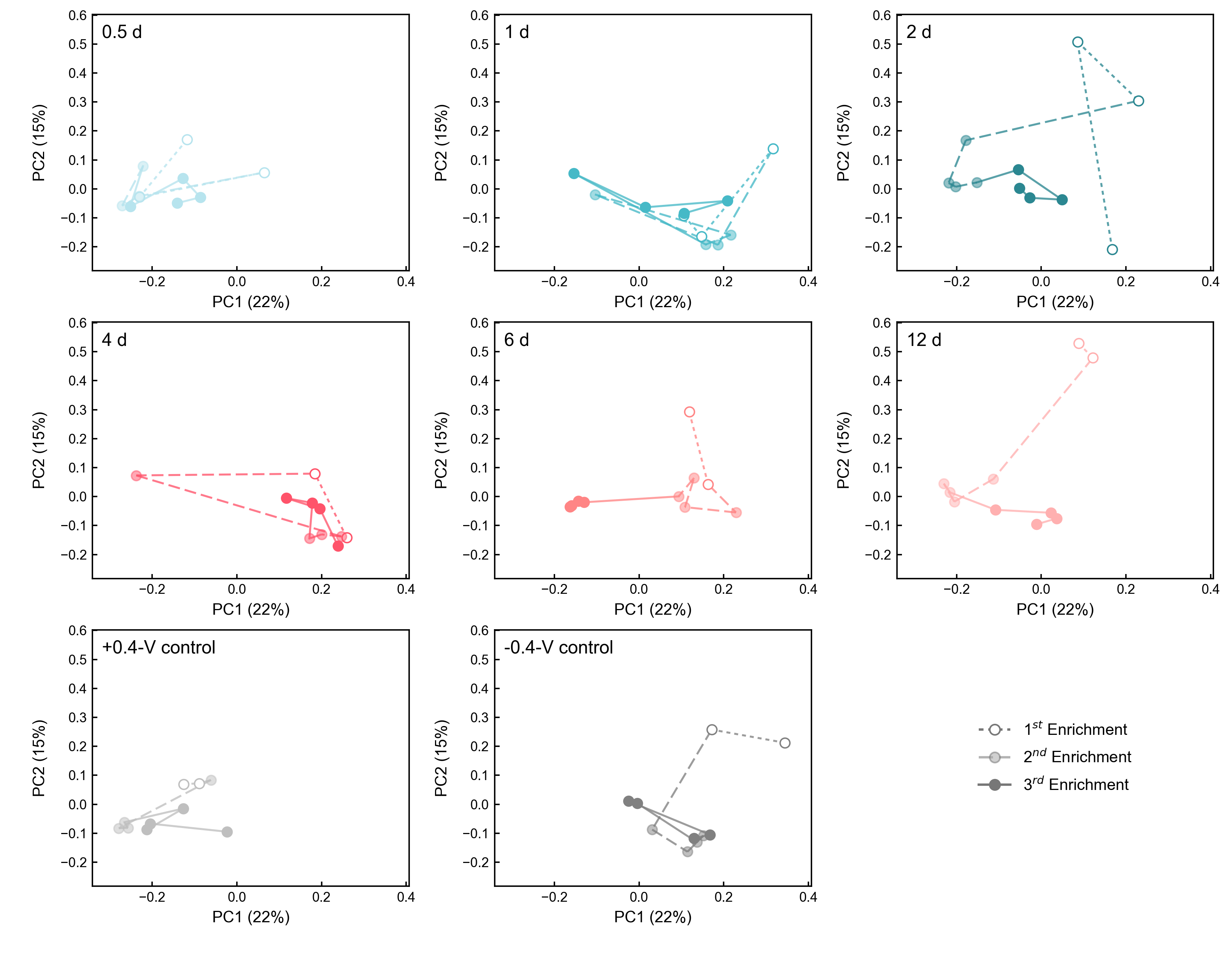

Figure S4. Bray-Curtis dissimilarity-based PCoA of community dynamics under eight conditions in three enrichments.

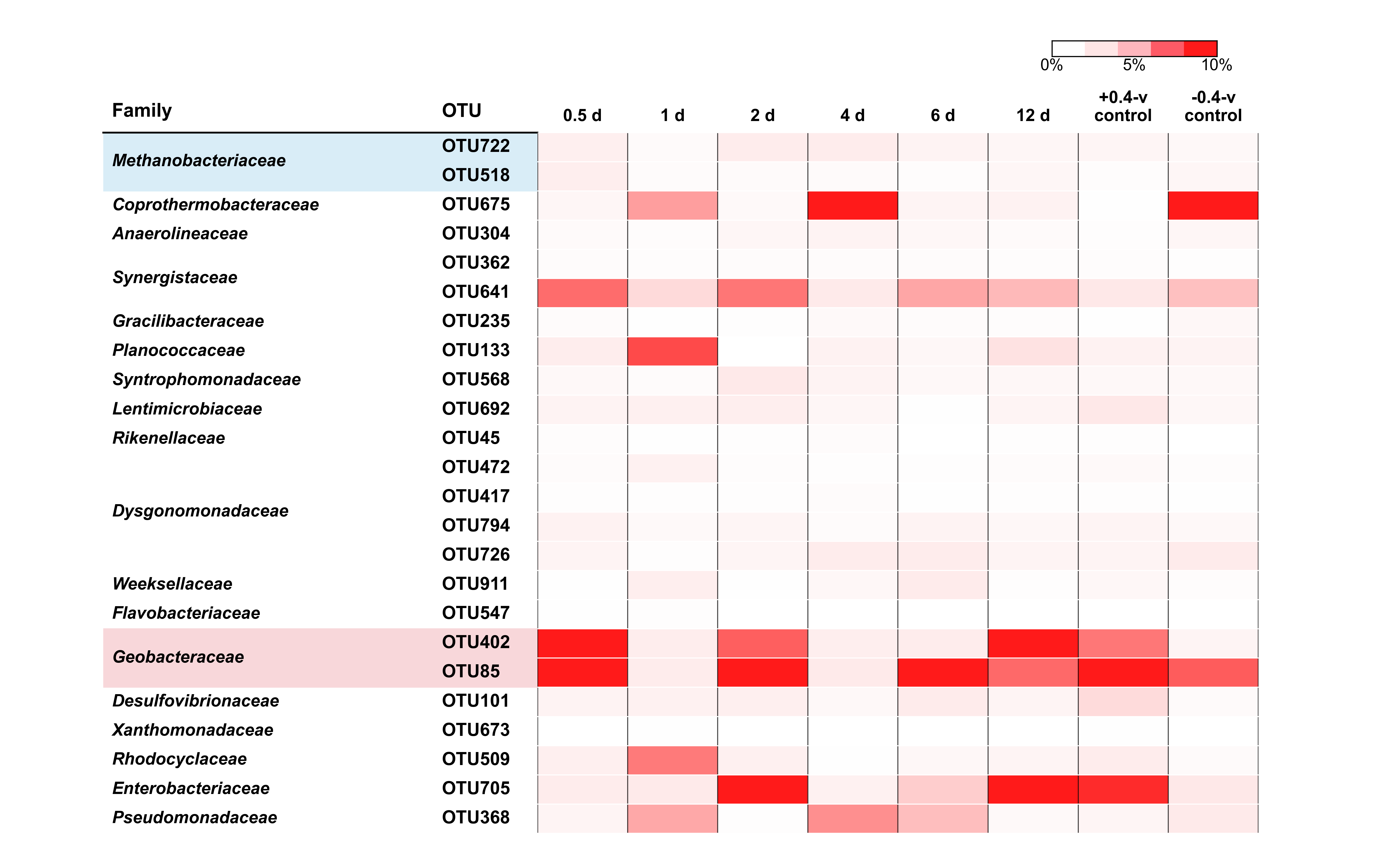

Figure S5. Activity (relative abundance of 16S rRNA transcript) of the 24 core populations under different alternating conditions.

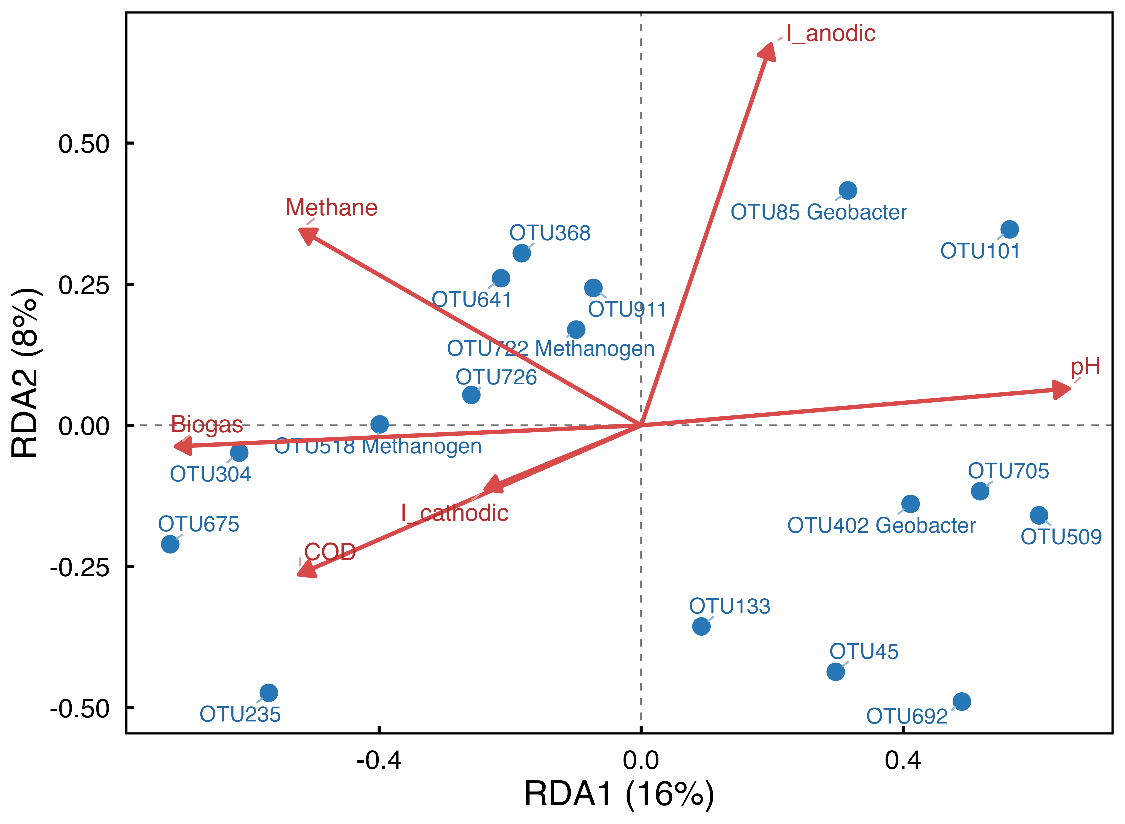

Figure S6. Redundancy analysis (RDA) of the relative growth activity of core populations derived from 16S rRNA transcript abundance in relation to environmental variables.

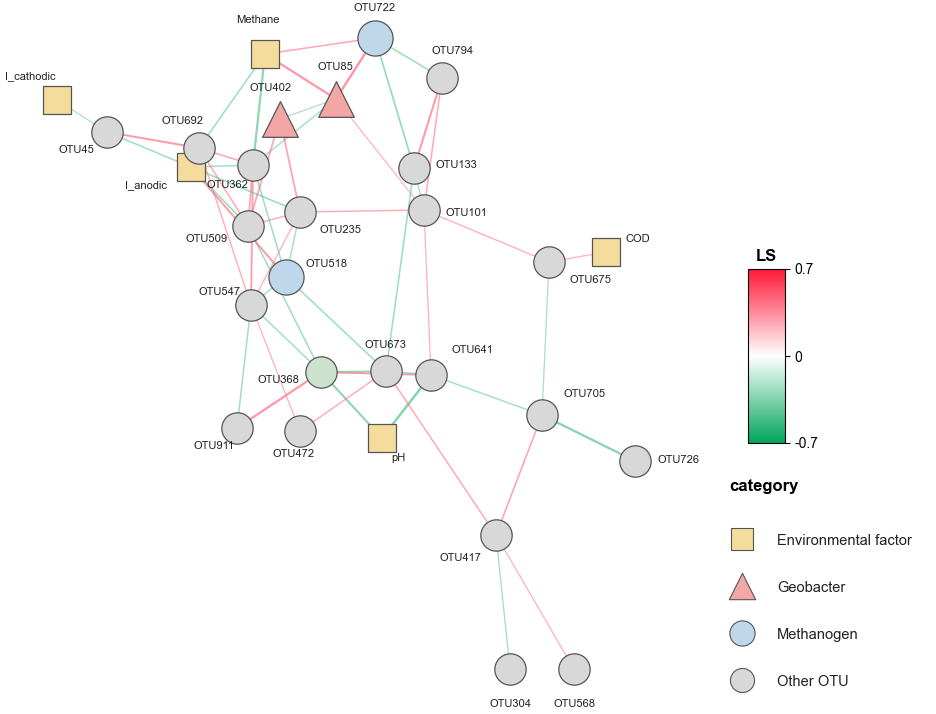

Figure S7. Integrated eLSA network of active core OTUs and environmental variables based on 16S rRNA transcript abundance
